# What determines the vertical structuring of pelagic ecosystems in the global ocean?

**DOI:** 10.1101/2024.07.04.602098

**Authors:** Mokrane Belharet, Matthieu Lengaigne, Nicolas Barrier, Andrew Brierley, Xabier Irigoien, Roland Proud, Olivier Maury

## Abstract

Offshore pelagic ecosystems are composed of vertically and functionally distinct epipelagic, migrant and resident mesopelagic communities. While this vertical structure plays a key role in carbon sequestration and in supporting important fisheries, there is still no consensus on the respective contribution of the environmental factors (light, oxygen) and processes controlling it at both global and regional scale. Here we combine mechanistic modelling and acoustic observations from the worldwide Malaspina scientific campaign to show that, while underwater light intensity is the primary factor controlling the vertical distribution and migration of pelagic organisms globally, oxygen plays a critical role in limiting the depth of migratory communities and the abundance of mesopelagic communities in Oxygen Minimum Zones. Furthermore, we show that a faithful reproduction of acoustic observations in some regions of the global ocean (southern Indian Ocean, western Pacific) cannot be achieved without separating migratory and resident mesopelagic communities into deep and shallow groups. By proposing a unified mechanistic model and an archetypical ecosystem structure constrained by comprehensive acoustic observations, this study provides a consistent understanding of the vertical structure and function of global pelagic ecosystems and paves the way for more reliable estimates of their climate-induced variability and change.

## Introduction

Offshore pelagic ecosystems are highly structured vertically (1, 2). The first maximum of animal biomass is generally located in the epipelagic zone, above 200m depth. This ocean-lit region is home to photosynthetic primary producers that fix atmospheric carbon and support highly productive and diverse fish populations, which are widely exploited by fisheries. Some of the organic carbon produced there is exported deeper in the water column by the biological pump (3, 4). A second biomass maximum is usually observed between 200m and 1000m depending on the region (5), in the mesopelagic zone which is too dark for photosynthesis but bright enough for some organisms to hunt by sight. Although largely untapped by fisheries and still poorly understood (6), the mesopelagic zone probably harbours the largest living biomass on the planet (7, 8) and plays an essential role in the biological carbon pump. Depending on the region, 20 to 90% of mesopelagic species (9) occupy it during the day, where they remain out of sight of epipelagic predators and come to the surface at night to feed on low trophic level epipelagic organisms. This massive diel vertical migration is the largest known animal migration on the planet (10).

This generic vertical structure is highly variable in space and time (11). Underwater light intensity and oxygen concentration have been identified as the key environmental variables controlling it (12–15). However, there is currently no consensus on their relative influence. While some studies conclude that oxygen controls the vertical distribution of fish, based on the correlation between their average vertical position and the oxygen content of the water column (9, 16), others argue that light is the dominant factor and exclude the direct effect of oxygen (17).

Here, we revisit this question by using a mechanistic approach based on the vertical component the APECOSM ecosystem model (18), which describes the diurnal and nocturnal vertical distribution of generic pelagic communities using an habitat-based advection diffusion equation driven by simple parameterizations of the physiological and behavioural responses of organisms to environmental factors (see method section and SI). We combine this model with low-frequency (38 kHz) echo sounder observations (i.e. a proxy for the vertical biomass density of gas-bladdered fish in the water column) from the Malaspina pan-oceanic scientific cruise (19, 20) to understand how light and oxygen structure pelagic ecosystems vertically. Specifically, we use these acoustic backscatter measures, that we correct using an acoustic model to account for the effects of body-size and vertical pressure on swimbladder resonance (see SI for further details), to estimate the parameters of a simple canonical model configuration driven by light intensity only. Then we identify the regions of the global ocean where the model deviates from the observations and we progressively complexify its structure and parameterization until we obtain a configuration that reproduces well the observations in all the ecological provinces crossed by the Malaspina cruise (Fig.1a). This allows us to propose a unified mechanistic framework to explain how pelagic ecosystems are structured vertically based on differential physiological and behavioural responses to light and oxygen.

**Fig.1:**
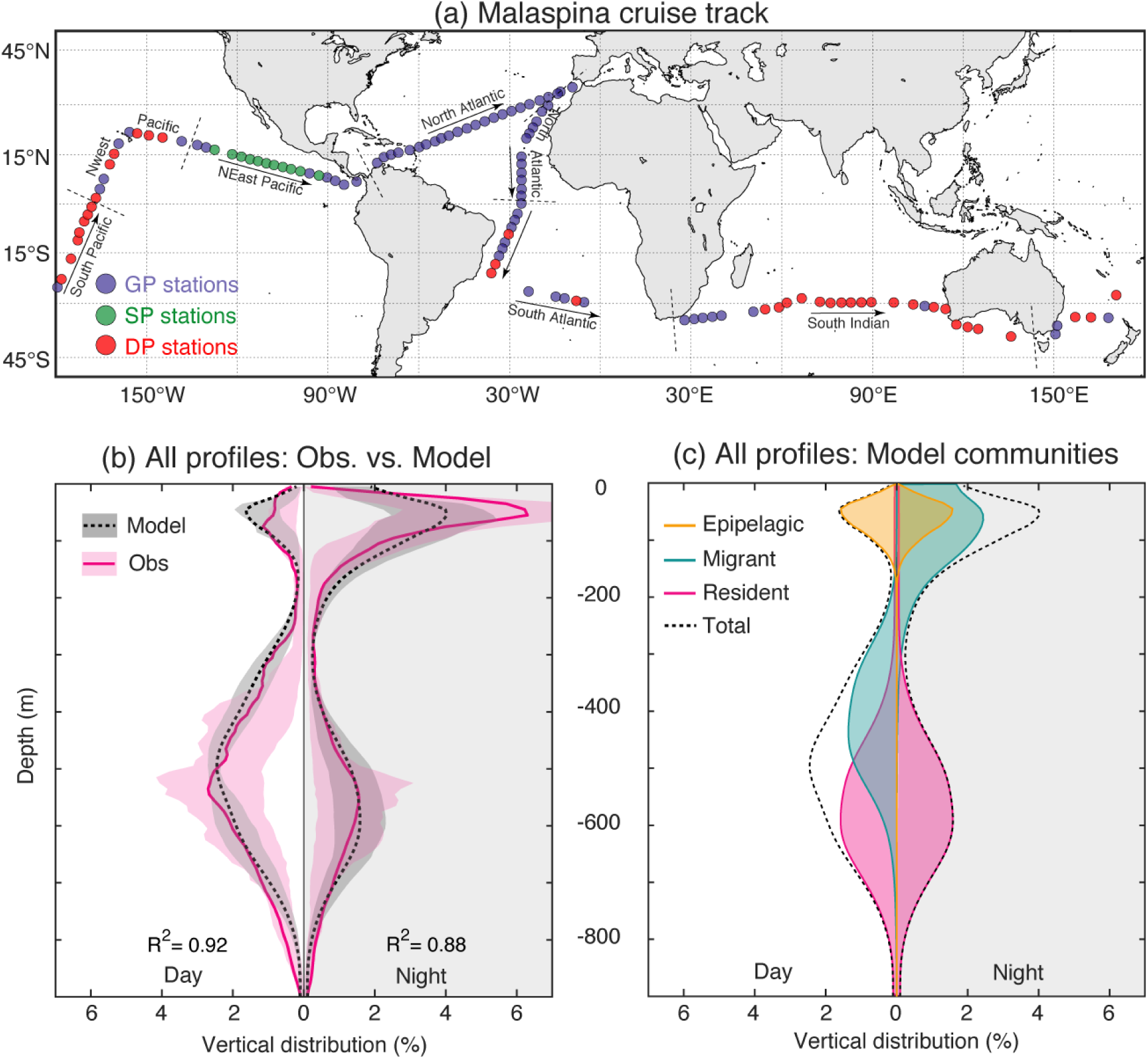
Global assessment of the three-communities light-based model. ***a***, Malaspina cruise track showing the location of the 122 acoustic stations used. Colours correspond to the groups of stations emerging in Fig. 2a and b. ***b***, Globally averaged daytime (left) and night-time (right) vertical profile for the 122 Malaspina stations (continuous line) and associated lower (25%) and upper (75%) quartiles (shading) for observations (pink) and model prediction (black). ***c***, Globally averaged daytime (left) and night-time (right) contribution of the three communities (epipelagic in yellow, migrant mesopelagic in cyan and resident mesopelagic in pink) to the total (dashed black) vertical profile in the simple three communities light-based model configuration.

## Results

### Light is the dominant factor at the global scale

Averaged over the entire Malaspina cruise (Fig. 1a), the observed backscattering acoustic profile (Fig. 1b) reveals the archetypical vertical structuring of pelagic ecosystems. The epipelagic biomass peak is more marked at night and the mesopelagic one, which lies at about 600m depth at night, grows and spreads up to 500m during the day. These observations, although globally consistent, do not shed light on the underlying community structure of the pelagic ecosystem, nor on its potential spatial variability. In an attempt to elucidate them, we first implement a simple configuration of the APECOSM model, which considers only three generic communities (one epipelagic, one resident mesopelagic and one migrant mesopelagic) as well as light as the only environmental forcing parameter, in order to assess its ability to simulate the vertical ecosystem structuring at the global scale.

As shown on Figure 1b, this simple configuration is already able to capture very satisfactorily the globally averaged vertical profile of acoustic data during daytime (R2 = 0.92) and night time (R2 = 0.88), with a very similar Deep Scattering Layer (DSL, i.e. the depth range where the acoustic signal is the strongest and where most mesopelagic biomass is concentrated) located between 400 and 700m. Thanks to the model, the average relative contributions of the different pelagic communities to the acoustic signal can be quantified and deconvoluted (Fig. 1c). On a global scale, the contribution of the epipelagic community, which occupies the surface layer night and day, is three times smaller than that of the migrating community. Similarly, the contribution of the migrating community to the acoustic signal is 20% smaller than that of the resident mesopelagic community, which is about 150m deeper on average during the day. The sum of the contributions of the three communities of this simple model makes it possible to predict the total acoustic signal, which matches well with the observations when averaged globally (Fig. 1b).

At the station level however, while the model accurately predicts the weighted mean depth (WMD) of the observed acoustic backscattering intensity for most of the night-time profiles (Fig.2a; R2 = 0.94), it performs worse for daytime (Fig. 2b; R2 = 0.71). This comes mainly from « Shallow Pattern » stations that are mostly located in the eastern tropical Pacific (SP, green dots on Fig.1a and 2b; 10% of profiles), where the predicted daytime WMDs are overestimated by more than 50m and from « Deep Pattern » stations in the Southern Indian Ocean and western Pacific (DP, red dots on Fig.1a and 2b; 31% of profiles), where daytime WMDs are underestimated by up to 100m. In all the other stations, that we call « Generic Pattern » (GP, blue dots on Fig.1a and 2b; 59% of profiles), predicted WMDs are consistent with observations, with an error being less than 50m.

**Fig.2 :**
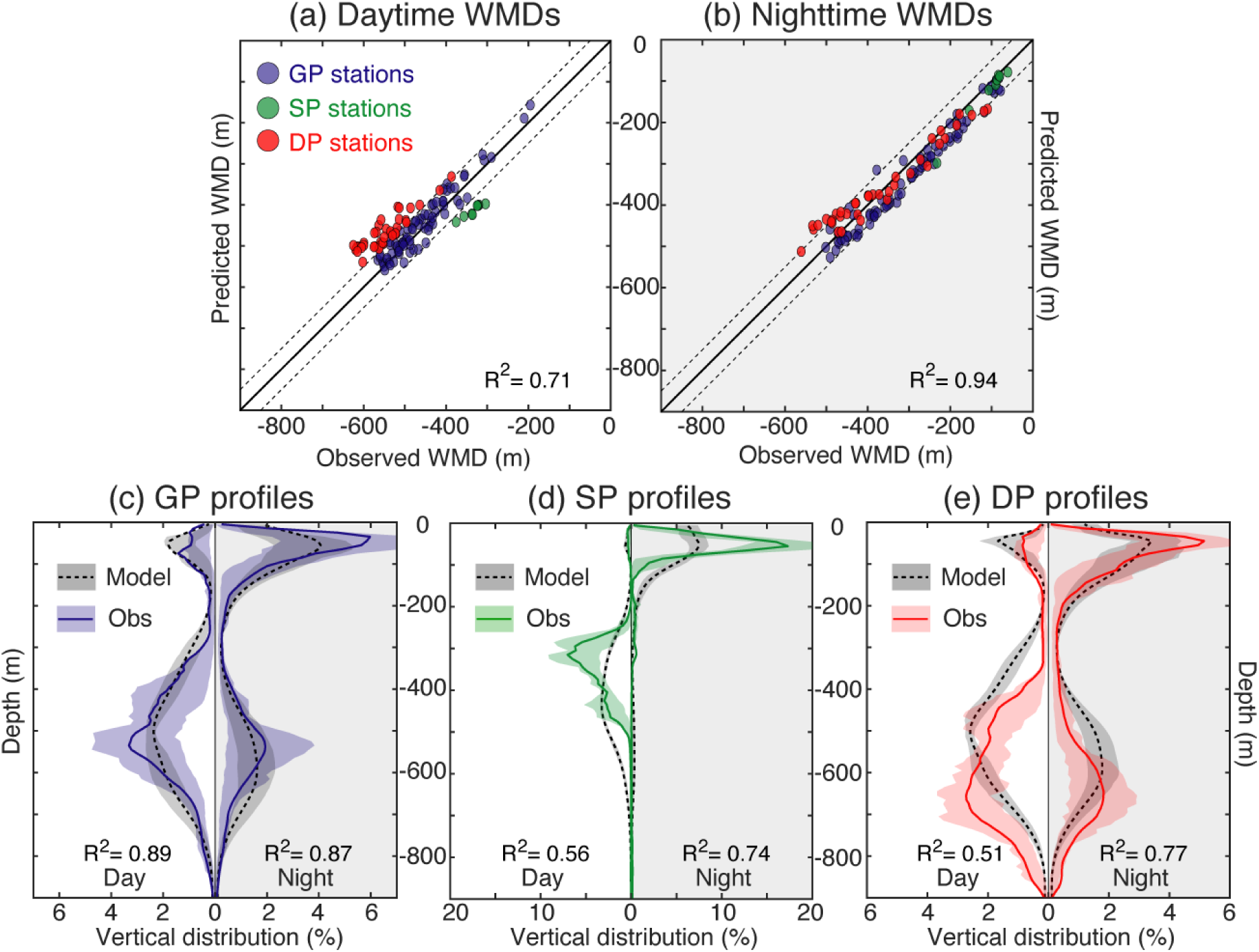
Regional biases of the three-communities light-based model. ***a***, Daytime and ***b***, night-time scatterplots of the predicted and observed WMDs (Weighted Mean Depths) for the 122 individual stations. Colours are assigned based on the difference between observed and predicted daytime WMDs (d = WMDobs - WMDmod): green if d ≤ -50m, red if d ≥ 50m and blue if -50m < d < 50m. ***c-e***, Predicted (black dashed lines) and observed (coloured lines) mean profiles and associated lower (25%) and upper (75%) quartiles (shadings) for the three groups (blue for GP, green for SP and red 122 for DP) during daytime (left) and night-time (right).

Fig. 2c-e evaluates the model’s ability to explain the observed 136 profiles for each of the three groups of stations identified in Fig. 2b. In accordance with the results in Fig. 2a-b, the model realistically simulates the averaged daytime (R2 = 0.89) and night-time (R2 = 0.87) vertical profiles of the GP stations (Fig. 2c), with a slight overestimation of the DSL depth for the deepest side of the profile. In contrast, the simulated averaged profiles are markedly biased in the SP and the DP groups (Fig. 2d-e), especially during the day where the agreement drops considerably (R2 = 0.56 and 0.51 respectively). These biases are largely associated with a diurnal DSL approximately 200m deeper in the model than in observations in the SP (Fig. 2d) and ∼100m shallower for the DP group (Fig. 2e). This indicates that our simple 3-communities light-based configuration of the model does not account for the key processes responsible for the specificities of the SP and DP groups.

### Oxygen limitation is key in the Oxygen Minimum Zones

As shown on Fig. 2d, the acoustic profiles in the eastern tropical Pacific (SP group) display a much narrower and shallower DSL than in other regions. Fig. 3a further indicates that the DSL is located at light intensity levels that are 100 to 1000 times higher in the SP group than for the two other groups, implying that even if underwater light intensity is slightly weaker in this region 17, light alone cannot explain the atypical DSL depth in the SP. The SP stations are located in a well-known Oxygen Minimum Zone (OMZ), characterized by a very steep and shallow oxycline (Fig. 3b) below which most organisms might not find enough oxygen to satisfy their metabolic needs. In addition to light, previous studies have therefore argued that oxygen is likely to play a key role in the particular vertical distribution of organisms in OMZs (14, 21, 22). When the response to dissolved oxygen concentration is included in the model (Fig.3c), the predicted diurnal DSL is indeed moving closer to the surface and becomes more pinched (Fig. 3c), while the predicted diurnal WMDs become much closer to observations (Fig. 3d). This striking improvement of the predicted daytime profiles (R2 increases from 0.56 to 0.90 when the effect of oxygen is included -Fig. 3c- and bias on WMD is reduced to less than 50m -Fig. 3d-) confirms that hypoxic waters limit the depth that can be attained by the migratory mesopelagic community during the day in the SP stations and prevents the resident mesopelagic organisms to live in the corresponding region (as will be shown later in Fig. 6). Thus, our model indicates that while migratory organisms can live in OMZs, their light-driven downward migration is halted at the oxycline depth at ∼0.7 ml.l-1 (Fig. S2), about 200m above their usual depth range (400-500 m) where light intensity is optimal but oxygen conditions unfavourable. This also explains why oxygen has no effect during the night when these migratory organisms occupy the epipelagic zone where oxygen levels are not limiting.

**Fig. 3:**
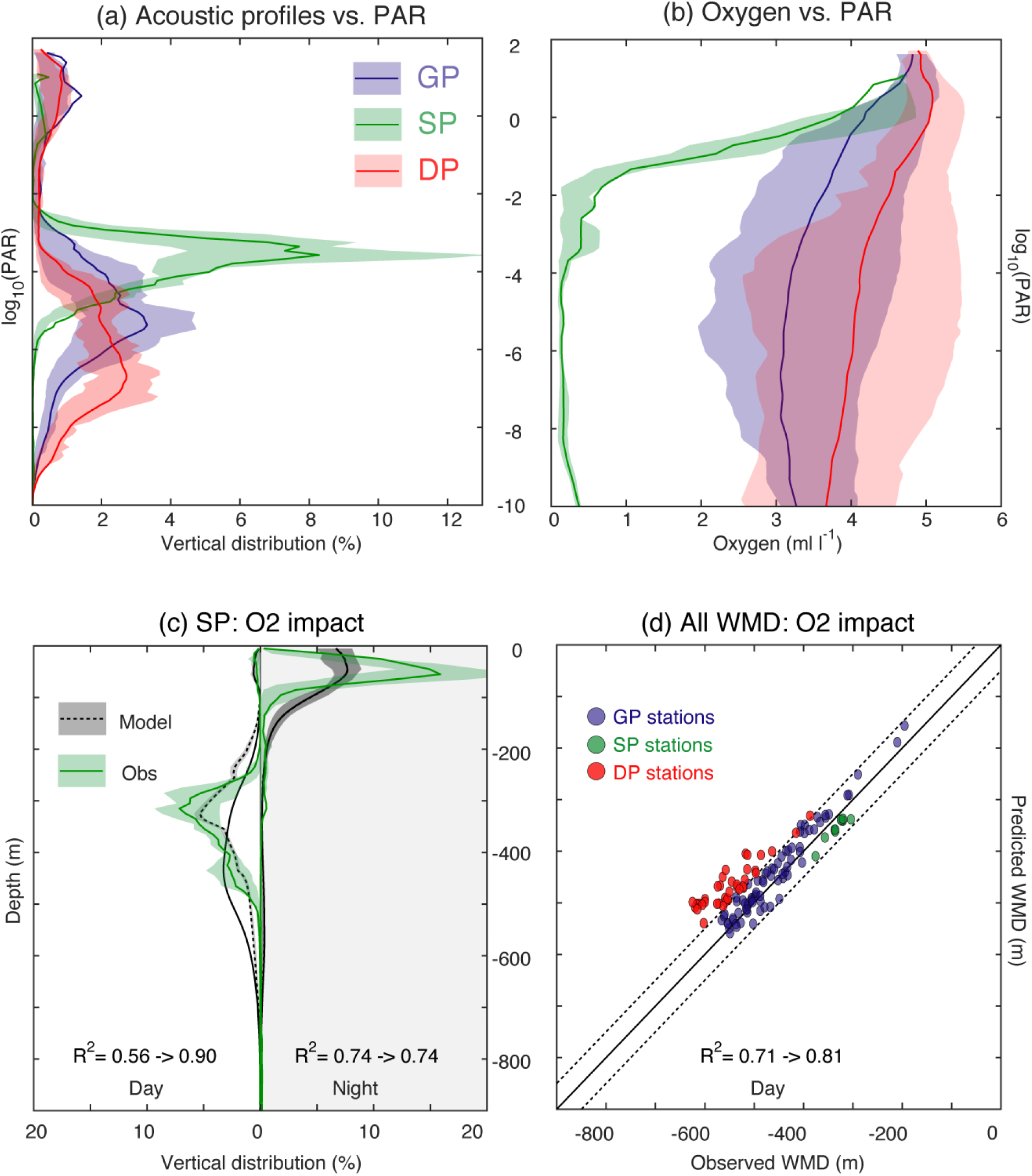
The role of oxygen in the Oxygen Minimum Zone. ***a,b***, Mean profiles of ***(a)*** acoustic backscatter data and ***(b)*** oxygen concentrations as a function of the decimal logarithm of observed light intensity (PAR) for the three groups of stations. ***c***, Predicted mean profiles in the SP group with (dashed black line) and without (continuous line) accounting for the effect of oxygen in the model compared to acoustic observations (green line). For ***a, b, c*** panels, shaded areas indicate lower (25%) and upper (75%) quartiles of the distribution. d, Scatter plot showing the correlation between predicted and observed daytime WMDs accounting for the effect of oxygen in the 3-community model.

### The regional importance of deep 185 mesopelagic communities

Although the inclusion of oxygen significantly improves the predicted profiles in the SP stations, this is not the case in the DP ones (Fig. 3d), where the model does not capture the anomalously deep observed DSL. This lack of oxygen limitation effect in the model is not surprising in the highly oxygenated DP waters that are very similar to that of the GP group (Fig. 3b). A detailed inspection of the echograms of the DP group reveals that they are not always homogenous and sometimes characterized by two distinct migratory and/or resident DSLs (see Fig. S6). In addition to the mesopelagic communities commonly observed between 300m and 600m, deeper mesopelagic communities are visible between 500 and 800m during the day, as reported in other regions (22–26). To assess whether this could explain the deeper DSL observed in the DP group, we added two deep mesopelagic communities (one migratory and one residents) to the model, in addition to the two previous shallow mesopelagic groups. Considering five communities instead of three and the effect of oxygen (what we call the « full model » configuration) significantly improves the agreement between predicted and observed WMDs at the global scale (Fig. 4a,b), solving in particular the problem of underestimation of WMDs during the day in the DP stations (Fig. 4a). As expected, the profile predicted by the full model is considerably improved for the DP group, with a much deeper simulated DSL than for the simple 3-community configuration, in closer agreement with observations (Fig. 4e).

**Fig.4 :**
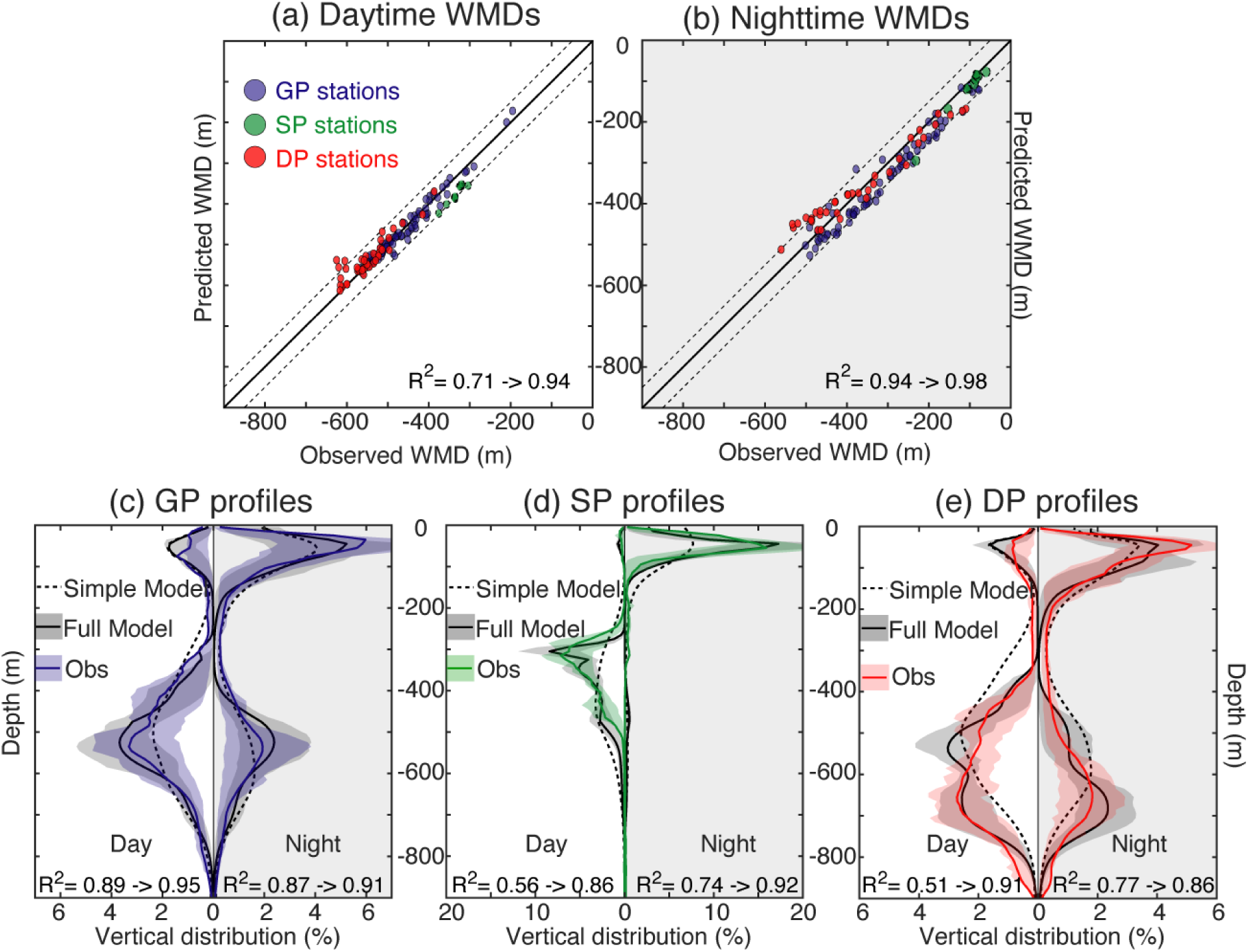
Including deep mesopelagic communities improves the model in DP. ***a***, Daytime and ***b***, night-time scatterplots between WMDs observed and predicted by the full model (including 5 communities and the effect of oxygen) for the 122 individual stations. ***c-e***, Mean profiles observed (coloured lines) and predicted by the full model (black lines) and associated lower (25%) and upper (75%) quartiles (shadings) for the three groups during daytime (left) and night-time (right). Mean profiles predicted by the simple model (3 communities and no oxygen effect) are reported as black dashed lines on each panel.

### Considering oxygen and deep communities improves the global model

In addition, including simultaneously the effects of oxygen and two deep communities not only improves the predicted vertical distributions in the DP and SP groups, but also in the GP group, both day and night (Fig. 4c,d). This is because even though the DP and SP regions are relatively marginal, de-biasing the model in these regions allows the global fit to be improved, including in the GP group.

While Fig. 4c-e allows the assessment of the model’s ability to reproduce average profiles in the three regions, Fig. 5a-c quantifies the model improvement at the individual station level. Moving from the simple to the full model improves the agreement between observed and predicted profiles, since both the median and the distribution of the coefficient R2 calculated for every individual profiles increases (Fig. 5a-c). The improvement is particularly clear in the SP (Fig. 5b) and in the DP (Fig. 5c), especially during the day, where 226 the median R2 and their distribution are very substantially improved (from 0.24 to 0.66 for SP and from 0.36 to 0.63 for DP). In these regions, R2 distributions between the simple and the full model overlap only marginally, indicating that most of the individual profiles have been improved by the complexification of the model and that the worst profiles in the full model are performing better than the best ones in the simple model. While the model is logically improved in those regions for which the addition of oxygen and deep communities were specifically done, it should be emphasized, as already pointed earlier (Fig. 4c), that it is also significantly improved in GP regions (the median R2 increased from 0.52 to 0.71) where the simple model was already performing relatively well (Fig. 4c and Fig. 5a).

**Fig. 5:**
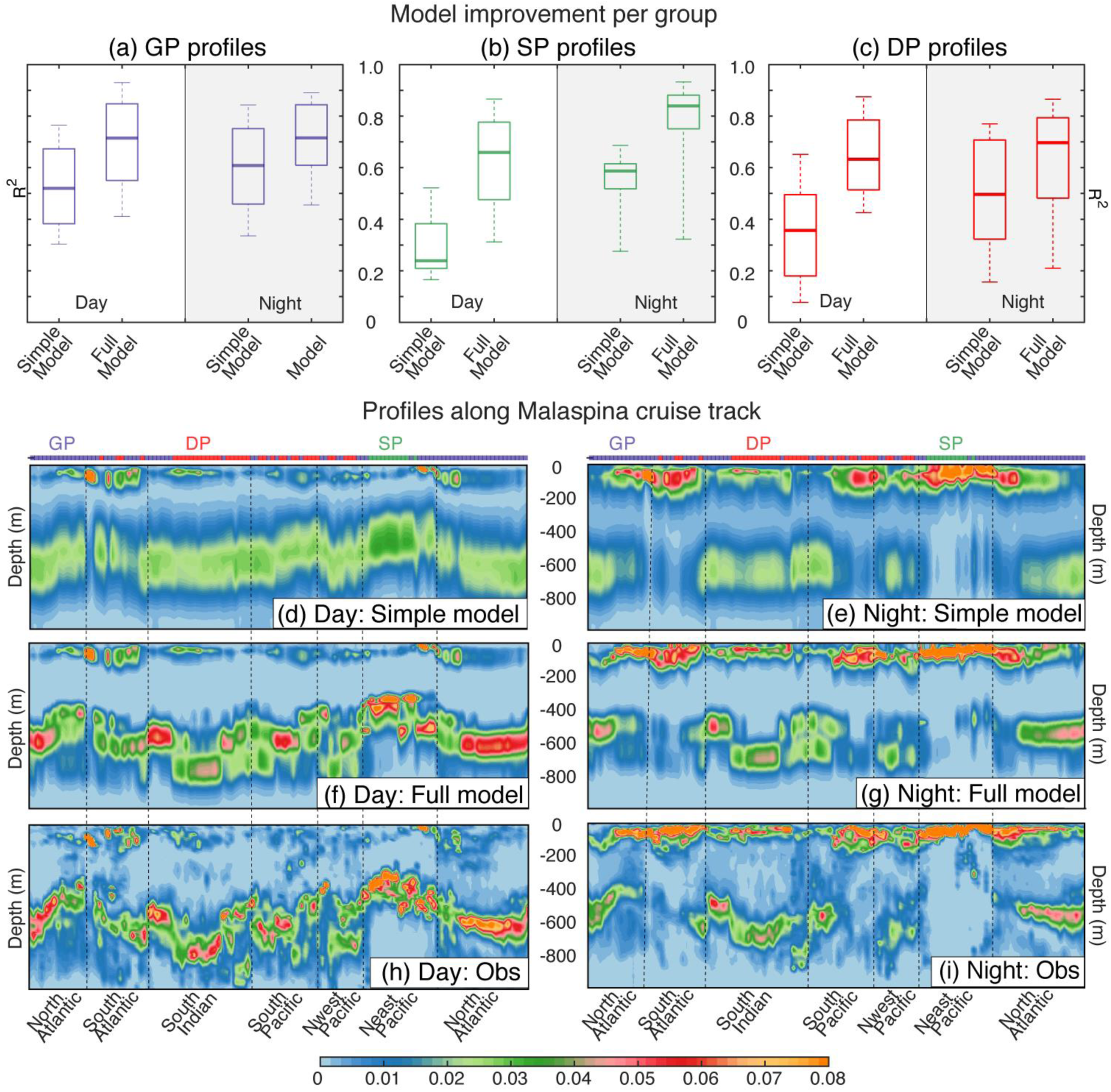
Synthetic evaluation of the simple and the full model configurations. ***a-c***, Boxplots comparing the two model configurations used in this study during the day (left) and the night (right) in the (***a***) General Pattern stations, (***b***) Shallow Pattern stations (***c***) Deep Pattern stations. The bold lines correspond to the median of the R2 calculated for all stations in the group, the box to the 25% and 75% quantiles and the whiskers to the 10% and 90% quartiles (***d-i***) Vertical distribution of the predicted and observed acoustic profiles along the Malaspina cruise route (cf. Fig. 1a) for the simple model configuration during daytime (***d***) and night-time (***e***), for the full model configuration during daytime (***f***) and night-time (***g***) and derived from acoustic observations during daytime (***h***) and nighttime (***i***). Dashed vertical lines correspond to the regional boundaries defined Fig. 1a.

This improvement between the simple and the full model is clearly seen in Fig. 5d-i when comparing the predictions from the two model configurations with the observations along the Malaspina transect. The full model (Fig. 5f,g) reproduces the observations (Fig. 5h,i) much more faithfully and in a much more contrasted way than the simple model configuration (Fig.5d,e). In particular, the fine structuring of the DSL and its geographical variability, which were poorly reproduced by the simple model, appear much finer and correspond very well to the observations, in all the oceanographically and ecologically very contrasted regions that were sampled by the Malaspina cruise, both day and night. The vertical distribution of the surface layer is also considerably improved at night.

### The model elucidates the global pelagic ecosystem structure and its regional variability

The full model elucidates the functional structure of global pelagic ecosystems by making it possible to estimate the relative contribution of each of the 5 communities to the total acoustic signal and its geographical variability 260 (Fig. 6a). Although heterogeneous and showing multiple and subtle geographical changes in dominance and association between the different groups, the structure of the GP group is in general dominated by shallow mesopelagic organisms (migrant and/or resident), with a lesser presence of epipelagic and deep mesopelagic organisms. On the contrary, the DP group is clearly dominated by migrant 265 and/or resident deep mesopelagic fish, to the detriment of the shallow mesopelagic groups, which almost totally disappear in the southern Indian Ocean. Finally, the SP group, essentially located in the eastern tropical Pacific OMZ, is exclusively dominated by shallow migrants which are extremely abundant there, in a totally atypical manner.

In addition to the community composition, the full model elucidates the vertical distribution and diel migrations of the different communities as well as their regional specificities. On average, the predicted relative community contributions for the GP group indicate a predominance of shallow communities for residents and a similar contribution of the deep and shallow migrants (Fig. 6b). In contrast in the SP stations, the average vertical distribution of biomass is mostly that of the shallow mesopelagic community that dominates the ecosystem and is displaced by the presence of the OMZ about 100m above its usual depth (Fig. 6c). Finally in the DP stations, deep migratory and resident mesopelagic communities predominate over shallow ones, resulting in a deep WMD (Fig. 6d) that is much more consistent with observations than when only two mesopelagic communities are considered (Fig. 4e).

**Fig. 6.**
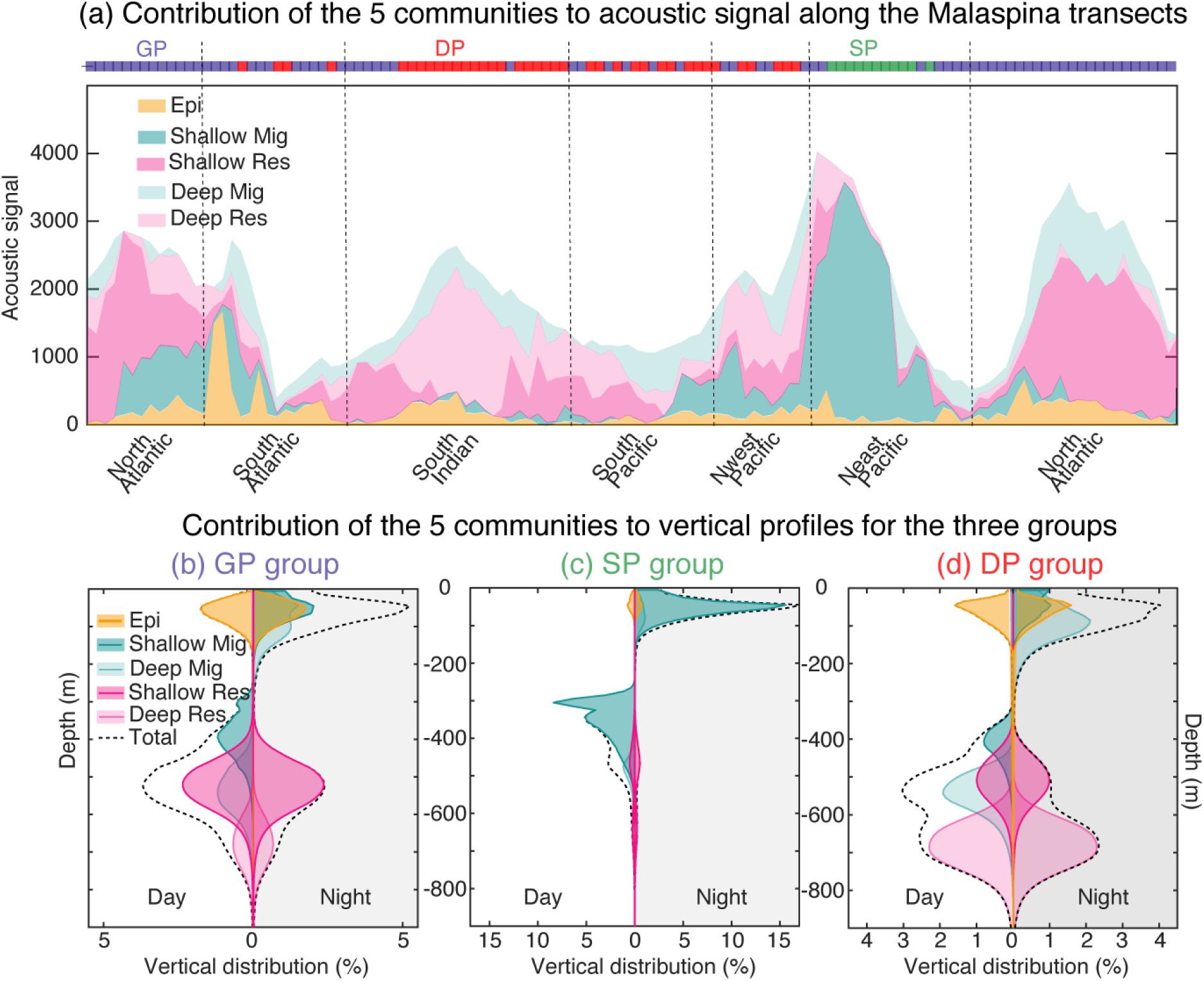
: Structure of oceanic pelagic ecosystems. (***a***) Contribution of the five pelagic communities to the acoustic signal predicted by the full model along the Malaspina transects. (***b-d***) Vertical acoustic profiles predicted by the full model for the five pelagic communities in the three groups identified. (***b***) General Pattern, (***c***) Shallow Pattern, (***d***) Deep Pattern

## Discussion

To our knowledge, this study is the first to examine the environmental factors responsible for the vertical distribution of pelagic communities using a mechanistic model applied to a dataset of acoustic backscatter profiles at global scale. This allows us to predict the diurnal and nocturnal relative biomass and vertical distribution of epipelagic and mesopelagic communities as a function of environmental drivers such as light intensity and dissolved oxygen. Our results first reveal that a simple model configuration explicitly representing three generic communities (epipelagic, migrant and resident mesopelagic), whose vertical habitat is influenced only by light intensity, is able to accurately reproduce the globally averaged profile of acoustic backscattering intensity (Fig. 1b). In agreement with previous studies (12, 13, 17, 27, 28), this indicates that light is the primary environmental factor governing the vertical distribution of the pelagic communities on a global scale and that different pelagic communities stay in distinct light intensity ranges.

Further to the predominant role of light, our results also reveal that oxygen plays a key role in the peculiar vertical structuring of organisms in the eastern Pacific OMZ. As in other regions with similar hypoxic conditions (29, 30), our acoustic data in the eastern Pacific display a much narrower and shallower DSL (peak in backscattering intensity at ∼320 m) than in well oxygenated regions (peak at ∼550m). We show that this layer is predominantly composed of shallow migratory organisms. Although few hypoxia-tolerant species have adapted their metabolism to temporarily or permanently inhabit OMZs with oxygen concentration below 0.1 ml l-1, such as the myctophid of the genus Diaphus observed in the quasi-anoxic waters of the Arabian Sea (29), the inclusion of dissolved oxygen in our mechanistic model demonstrates that the extremely low oxygen concentrations in the middle of the water column in the eastern tropical Pacific are not suitable for most mesopelagic organisms and are actually responsible for the shallow DSL for SP profiles. This is consistent with in situ observations (14, 21) and indicates that deoxygenated waters form a barrier that prevents migrating organisms from reaching depths where light conditions are generally optimal and precludes the presence of a resident community. In the OMZ region, migrating organisms are indeed located between 235 and 315 m depth where oxygen conditions are around 0.5 ml l-1 and are much less present deeper where oxygen drops well below this threshold (Fig. 3a-c). These findings are at odds with those of Aksnes et al. (2017), which suggests that high concentrations of the coloured dissolved organic matter (CDOM, which is an important light absorber) lead to a reduction in light penetration in the mesopelagic zone of OMZ that would be sufficient to explain the shallow distribution of DSL in this region, thus ruling out the potential direct effect of oxygen. In line with our results (Fig. 2a), the diurnal distribution of acoustic backscatter in Aksnes et al. (2017) (their Fig. 2c ) clearly shows that the DSL of the low oxygen stations is located in waters with much higher irradiance (> 2 orders of magnitude) than those of the high oxygen stations. Thus, the high CDOM concentrations and the resulting reduced irradiance in this region cannot alone explain the shallow location of DSL.

Ecologically, since most migrating organisms are blocked above the OMZ, where oxygen concentrations are low but still tolerable (between 0.1 and 0.7 ml l^-1^), they are forced to stay in brighter waters than usual and might experience higher risks of visual predation. However, most observations suggest that the moderate hypoxic conditions in the top of the OMZ where migratory mesopelagics stay might still provide a refuge from large epipelagic visual predators, which have higher oxygen requirements than mesopelagic organisms (13, 14, 26, 31, 32). Migrating species in OMZ regions indeed use a variety of adaptations to cope with moderate hypoxia, such as reduced activity or anaerobic metabolism during the day with migrations to oxygen-rich waters close to the surface at night to re-oxygenate (33–36). On the contrary, resident organisms that cannot migrate upwards have no possibility to adapt to hypoxia, which explains their very low relative biomass estimated by our model in the eastern tropical Pacific OMZ (Fig. 6d). In contrast to the GP and SP stations, the echograms of the DP group show two more or less visible vertically migrating DSLs. This specific vertical structuring has already been observed in other oceanic regions (9), such as in the North Atlantic gyre (23–25), the north western part of the eastern Central Pacific (Clarion-Clipperton zone (22)), the California coast (36) and the Red Sea (26). In addition to the shallow migrant and resident mesopelagic communities, the inclusion of deep mesopelagic communities in our model allows us to reproduce the vertical structuring observed regionally across the sampled regions (cf. Fig.4e). This leads to a significant improvement over the three-community configuration (Fig. 5c) and to a more faithful reproduction of the echograms collected throughout the Malaspina cruise (Fig. 5f-i). Although the taxonomic composition of the pelagic communities is not available empirically, our model suggests that the structure (i.e. the community composition) of the pelagic ecosystem varies considerably geographically. These structural changes that our model captures provide a mechanistic explanation for the 6 homogeneous “echobiomes” that have been identified from global acoustic observations (37) and that correspond quite well to the mesopelagic biogeochemical provinces previously defined on the basis of ocean physicochemical properties (5).

Finally, although this study is based only on data from the Malaspina campaign, along a single transect between 40°S and 40°N latitude, our mechanistic approach allows predicting the vertical distributions of the five communities under consideration at any point in the ocean on a global scale (see Fig. S5 in SI). It could therefore be used to explore potential ecosystem responses to projected evolution of environmental conditions induced by climate change, such as water warming and increasing stratification, deoxygenation and expansion of OMZs, and potential changes in light conditions due to changes in cloud cover and water turbidity (38). Such projections were previously performed with the full APECOSM model (38, 39) using the simple 3-community configuration without data-driven parameter estimation. The present study paves the way for more realistic projections based on our full model configuration and the better understanding of the processes involved it underlies.

## Materials and Methods

### Data description

#### Acoustic data

In this study we used acoustic backscattering intensity data collected using a Simrad EK60 echosounder operating at 38-kHz as a proxy for fish biomass. These data were collected during the Malaspina Circumnavigation Expedition 20 that was conducted between December 2010 and July 2011 through the tropical and subtropical Atlantic, Indian and Pacific Oceans. We used a total of 122 x 2 vertical acoustic profiles collected at different stations (Fig 1.a) where both nocturnal and diurnal profiles were available. Data were cleaned (noise spikes and attenuated signal were removed), plotted and visually checked to assess data quality; in some instances, data were removed where false bottom detections were made. Each profile extends from 0 to 1000m depth with a vertical resolution of 10m. Day and night were distinguished on the basis of the sun angle information, with day when the sun angle is higher than 0° and night when it is lower than -18°.

#### Environmental data

Our model of the vertical distribution of ecological communities is driven by light (approximated by the Photosynthetically Active Radiation -PAR-), temperature and dissolved oxygen concentration. The underwater irradiance observed during the Malaspina cruise was measured accurately to a minimum level of 0.03 µmol quanta m^-2^ s^-1^, which corresponded to the daytime irradiance at depths ranging from 150 to 280 m depending on the water clarity (17). Therefore, these vertical profiles only cover the epipelagic water column, whereas in this study we aim to consider both epipelagic and mesopelagic realms (0 - 1000 m depth). To overcome this limitation, we used the vertical profiles of PAR simulated with the coupled Physics-Biogeochemistry ocean model NEMO-PISCES (40) forced over the 1958-2018 period with atmospheric inputs from the JRA atmospheric reanalysis (41), which is representative of surface atmospheric variability observed over the historical period (42). We have averaged the monthly PAR outputs over 8 years (2010–2018) to derive a monthly climatology representative of the Malaspina years. Despite limited differences in the OMZ regions (<50m), the simulated vertical profiles of PAR are very consistent with the observed ones (Malaspina) over the entire epipelagic water column (Fig. S7), above the abrupt decrease that appears at the end of the observed profiles, and which results from an observational bias related to the sensor sensitivity (17).

The seawater dissolved oxygen and temperature were obtained from the World Ocean Atlas 2013 (43), Environmental data are provided on 3D grids with a 1° horizontal resolution and 75 vertical levels with an increased resolution near the surface. The environmental drivers were interpolated linearly at the location of the 122 acoustic stations from their original grids, taking into account the geographic coordinates of the stations, their sampling dates and the 10m vertical discretization of the acoustic profiles.

### Model Description

#### Vertical movement

In this study, we use the component of the Apex Predators ECOsystem Model APECOSM (18) that represents the vertical movements and distribution of pelagic communities. In APECOSM, vertical movements are supposed to obey the following habitat-based advection-diffusion equation

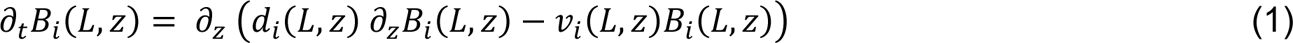

With *B_i_*(*L*, *z*) being the biomass density of community *i* organisms of scaled length *L* at depth *z*. The vertical velocity *v_i_*(*L*, *z*) is assumed to be proportional to the gradient of the habitat function 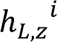 that characterizes the quality of the habitat for community *i* . It is also assumed to increase proportionally to body length and to decrease linearly with increasing habitat quality to ensure that organisms swim toward better habitats when they are in poor habitats but don’t look for good habitats when they have reached them:

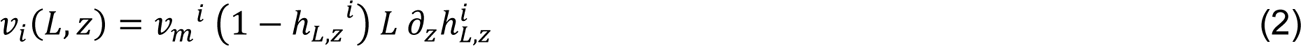

The vertical diffusion coefficient *d_z_* is supposed to have two components. The first one, which represents random foraging vertical movements, is proportional to the squared scaled length of organisms and it increases linearly with the habitat function 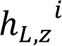 so that organisms spend more time randomly looking for food when their habitat is good. The second component is a constant diffusion term 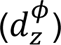 accounting for purely physical vertical mixing. It is especially important for small organisms:

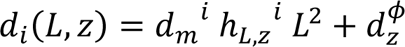

#### Habitat

The vertical habitat function 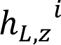 quantifies the functional response of individuals to their environment. Here, we simply assume that it is equal to the product of independent response functions to light, oxygen and temperature:

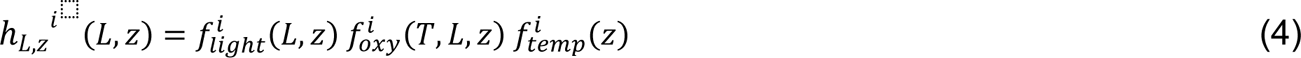

where, 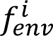 is the scaled response function of the generic community *i* to environmental factors *enc* = {*light,oxy,temp*}. Various physiological and behavioral processes are implicit in the definition of 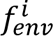 functions. These are described in detail in the SI.

The vertical distribution of all ecological communities considered here is governed by equation (1) that is integrated analytically. Each community is distinguished from the others by the value of movement and habitat parameters that capture the particularities of their vertical behaviors as detailed below.

#### Generic communities

Open ocean pelagic organisms are divided into three generic communities based on their position in the water column during the day and whether or not they are capable of diel vertical migration: *(i)* the epipelagic community (*EC*), which occupies the euphotic zone (from the surface to about 200 m depth); *(ii)* the Resident mesopelagic community (*RMC*), which occupies the twilight zone (200 – 1000 m); *(iii)* the Migrating mesopelagic community (*MMC*), which occupies the twilight zone during the day and migrates to the surface during the night. For the purpose of this study, we further divided the migratory and resident mesopelagic communities into two sub-communities: shallow (MMC_S_ and RMC_S_) and deep (MMC_D_ and RMC_D_).

#### Vertical community profiles

Unlike the acoustic signal, which is undifferentiated, our model distinguishes different ecological communities and provides estimates of their relative biomasses and vertical distribution. To represent the vertical distribution of the different communities, we calculate their vertical profile, the vertical density distribution of their biomass:

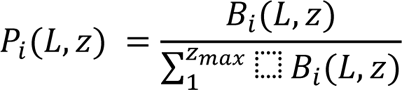

where *B_i_ (L, z)* is the biomass of organisms of length *L* belonging to the generic community *i* at depth *z* which is obtained by integrating equation (1), and *z_max_* is the total number of depth levels considered.

The aggregated biomass profile combining all the communities corresponds to the sum of the community profiles weighted by their relative biomass:

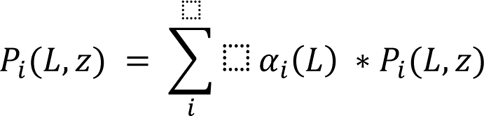

where *i = {EC, MMC , MMC , RMC , RMC }*, and 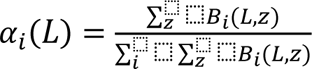 is the relative biomass of community *i* at length *L* compared to the total biomass of all communities at the same length.

### Model Parameters and Calibration

We estimated the model parameters by computing the minimum log-likelihood of the acoustic observations using the AD model builder (ADMB) software (44) (see SI for more details).

For that purpose, we assumed a Gaussian distribution of errors between the community-aggregated vertical profiles *P_mod_* predicted by the model and the observed acoustic profiles *P_obs_*. Considering the different stations (s) where the acoustic data were collected and both day and night periods (*η*), we therefore minimized the objective function

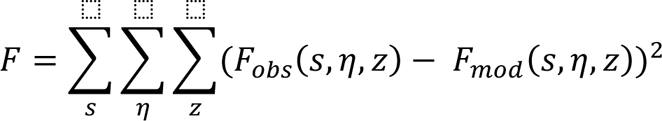

where,

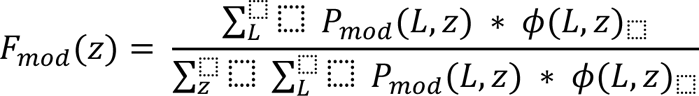

with *ϕ*(*L*, *z*) a correction factor implemented to account for the nonlinear distortion of the acoustic signal with depth and organisms size (see S3 in SI for more details), and *F_obs_* the observed vertical profile of the acoustic backscatter *P_obs_* normalized over the water column (between 0 and 1000 m depth).

## Acknowledgements

The present work has been carried out in the framework of the SUMMER project, funded by the European Union’s Horizon 2020 research and innovation programme under grant agreement No 817806.

## References

1. R. Proud, M. Cox, C. Le Guen, A. Brierley, Fine-scale depth structure of pelagic communities throughout the global ocean based on acoustic sound scattering layers. Mar. Ecol. Prog. Ser. 598, 35–48 (2018).

2. R. Proud, M. J. Cox, A. S. Brierley, Biogeography of the Global Ocean’s Mesopelagic Zone. Curr. Biol. 27, 113–119 (2017).

3. A. Martin, et al., Study the twilight zone before it is too late. 3 (2020).

4. D. A. Siegel, T. DeVries, I. Cetinić, K. M. Bisson, Quantifying the Ocean’s Biological Pump and Its Carbon Cycle Impacts on Global Scales. Annu. Rev. Mar. Sci. 15, annurev-marine-040722–115226 (2023).

5. G. Reygondeau, et al., Global biogeochemical provinces of the mesopelagic zone. J. Biogeogr. 45, 500–514 (2018).

6. M. A. St. John, et al., A Dark Hole in Our Understanding of Marine Ecosystems and Their Services: Perspectives from the Mesopelagic Community. Front. Mar. Sci. 3 (2016).

7. X. Irigoien, et al., Large mesopelagic fishes biomass and trophic efficiency in the open ocean. Nat. Commun. 5, 3271 (2014).

8. R. Proud, N. O. Handegard, R. J. Kloser, M. J. Cox, A. S. Brierley, From siphonophores to deep scattering layers: uncertainty ranges for the estimation of global mesopelagic fish biomass. ICES J. Mar. Sci. 76, 718–733 (2019).

9. T. A. Klevjer, et al., Large scale patterns in vertical distribution and behaviour of mesopelagic scattering layers. Sci. Rep. 6, 19873 (2016).

10. G. C. Hays, A review of the adaptive significance and ecosystem consequences of zooplankton diel vertical migrations. Hydrobiologia 503, 163–170 (2003).

11. T. T. Sutton, et al., A global biogeographic classification of the mesopelagic zone. Deep Sea Res. Part Oceanogr. Res. Pap. 126, 85–102 (2017).

12. A. Røstad, S. Kaartvedt, D. L. Aksnes, Light comfort zones of mesopelagic acoustic scattering layers in two contrasting optical environments. Deep Sea Res. Part Oceanogr. Res. Pap. 113, 1–6 (2016).

13. T. Langbehn, D. Aksnes, S. Kaartvedt, Ø. Fiksen, C. Jørgensen, Light comfort zone in a mesopelagic fish emerges from adaptive behaviour along a latitudinal gradient. Mar. Ecol. Prog. Ser. 623, 161–174 (2019).

14. A. N. Netburn, J. Anthony Koslow, Dissolved oxygen as a constraint on daytime deep scattering layer depth in the southern California current ecosystem. Deep Sea Res. Part Oceanogr. Res. Pap. 104, 149–158 (2015).

15. A. E. Maas, S. L. Frazar, D. M. Outram, B. A. Seibel, K. F. Wishner, Fine-scale vertical distribution of macroplankton and micronekton in the Eastern Tropical North Pacific in association with an oxygen minimum zone. J. Plankton Res. 36, 1557–1575 (2014).

16. D. Bianchi, E. D. Galbraith, D. A. Carozza, K. A. S. Mislan, C. A. Stock, Intensification of open-ocean oxygen depletion by vertically migrating animals. Nat. Geosci. 6, 545– 548 (2013).

17. D. L. Aksnes, et al., Light penetration structures the deep acoustic scattering layers in the global ocean. Sci. Adv. 3, e1602468 (2017).

18. O. Maury, An overview of APECOSM, a spatialized mass balanced “Apex Predators ECOSystem Model” to study physiologically structured tuna population dynamics in their ecosystem. Prog. Oceanogr. 84, 113–117 (2010).

19. C. M. Duarte, Seafaring in the 21St Century: The Malaspina 2010 Circumnavigation Expedition. Limnol. Oceanogr. Bull. 24, 11–14 (2015).

20. Martinez, Udane, et al., Raw EK60 echosounder data (38 and 120 kHz) collected during the Malaspina 2010 Spanish Circumnavigation Expedition (14th December 2010, Cádiz - 14th July 2011, Cartagena). PANGAEA - Data Publisher for Earth & Environmental Science. 10.1594/PANGAEA.921760. Deposited 2020.

21. W. Ekau, H. Auel, H.-O. Pörtner, D. Gilbert, Impacts of hypoxia on the structure and processes in pelagic communities (zooplankton, macro-invertebrates and fish). Biogeosciences 7, 1669–1699 (2010).

22. J. N. Perelman, E. Firing, J. M. A. van der Grient, B. A. Jones, J. C. Drazen, Mesopelagic Scattering Layer Behaviors Across the Clarion-Clipperton Zone: Implications for Deep-Sea Mining. Front. Mar. Sci. 8, 632764 (2021).

23. A. Ariza, et al., Vertical distribution, composition and migratory patterns of acoustic scattering layers in the Canary Islands. J. Mar. Syst. 157, 82–91 (2016).

24. L. Marohn, et al., Distribution and diel vertical migration of mesopelagic fishes in the Southern Sargasso Sea — observations through hydroacoustics and stratified catches. Mar. Biodivers. 51, 87 (2021).

25. T. T. Sutton, Vertical ecology of the pelagic ocean: classical patterns and new perspectives: vertical ecology of the pelagic ocean. J. Fish Biol. 83, 1508–1527 (2013).

26. E. Dypvik, T. A. Klevjer, S. Kaartvedt, Inverse vertical migration and feeding in glacier lanternfish (Benthosema glaciale). Mar. Biol. 159, 443–453 (2012).

27. T. A. Klevjer, W. Melle, T. Knutsen, D. L. Aksnes, Vertical distribution and migration of mesopelagic scatterers in four north Atlantic basins. Deep Sea Res. Part II Top. Stud. Oceanogr. 180, 104811 (2020).

28. N. Dupont, T. A. Klevjer, S. Kaartvedt, D. L. Aksnesa, Diel vertical migration of the deep-water jellyfish Periphylla periphylla simulated as individual responses to absolute light intensity. Limnol. Oceanogr. 54, 1765–1775 (2009).

29. J. Kinzer, R. Bottgerschnack, K. Schulz, Aspects of Horizontal Distribution and Diet of Myctophid Fish in the Arabian Sea with Reference to the Deep-Water Oxygen Deficiency. Deep-Sea Res. Part Ii-Top. Stud. Oceanogr. 40, 783–800 (1993).

30. J. Gjøsæter, Abundance and production of lanternfish (Myctophidae) in the western and northern Arabian Sea. (1981).

31. A. C. Lavery, D. Chu, J. N. Moum, Measurements of acoustic scattering from zooplankton and oceanic microstructure using a broadband echosounder. ICES J. Mar. Sci. 67, 379–394 (2010).

32. A. Bertrand, M. A. Barbieri, F. Gerlotto, F. Leiva, J. Córdova, Determinism and plasticity of fish schooling behaviour as exemplified by the South Pacific jack mackerel Trachurus murphyi. Mar. Ecol. Prog. Ser. 311, 145–156 (2006).

33. B. A. Seibel, Critical oxygen levels and metabolic suppression in oceanic oxygen minimum zones. J. Exp. Biol. 214, 326–336 (2011).

34. L. Stramma, et al., Expansion of oxygen minimum zones may reduce available habitat for tropical pelagic fishes. Nat. Clim. Change 2, 33–37 (2012).

35. J. J. Childress, B. A. Seibel, Life at stable low oxygen levels: adaptations of animals to oceanic oxygen minimum layers. J. Exp. Biol. 201, 1223–1232 (1998).

36. S. S. Urmy, J. K. Horne, D. H. Barbee, Measuring the vertical distributional variability of pelagic fauna in Monterey Bay. ICES J. Mar. Sci. 69, 184–196 (2012).

37. A. Ariza, et al., Global decline of pelagic fauna in a warmer ocean. Nat. Clim. Change 12, 928–934 (2022).

38. D. P. Tittensor, et al., Next-generation ensemble projections reveal higher climate risks for marine ecosystems. Nat. Clim. Change 11, 973–981 (2021).

39. H. K. Lotze, et al., Global ensemble projections reveal trophic amplification of ocean biomass declines with climate change. Proc. Natl. Acad. Sci. 116, 12907–12912 (2019).

40. O. Aumont, C. Ethé, A. Tagliabue, L. Bopp, M. Gehlen, PISCES-v2: an ocean biogeochemical model for carbon and ecosystem studies. Geosci. Model Dev. 8, 2465– 2513 (2015).

41. S. Kobayashi, et al., The JRA-55 Reanalysis: General Specifications and Basic Characteristics. J. Meteorol. Soc. Jpn. Ser II 93, 5–48 (2015).

42. N. Barrier, et al., Mechanisms underlying the epipelagic ecosystem response to ENSO in the equatorial Pacific ocean. Prog. Oceanogr. 213, 103002 (2023).

43. S. Levitus, et al., World Ocean Atlas 2013 (NCEI Accession 0114815). NOAA National Centers for Environmental Information. 10.7289/V5F769GT. Deposited 2015.

44. D. A. Fournier, et al., AD Model Builder: using automatic differentiation for statistical inference of highly parameterized complex nonlinear models. Optim. Methods Softw. 27, 233–249 (2012).

