## Supplementary Information for "What determines the vertical structuring of pelagic ecosystems in the global ocean?"

### 1) Functional responses to environmental variables

**Functional Response to Light Intensity**

Light intensity is an important factor governing the vertical distribution of pelagic organisms in the water column. Concretely, pelagic communities are believed to have a preference for a range of light intensity referred to as the Light Comfort Zone (LCZ) (1–4), which plays an important role in defining the daytime depth position and migration range of mesopelagic communities.

The sensitivity of vision to light is assumed to be a Gaussian function of the logarithm of light (5, 6) *i.e.* a lognormal function of light and can be written as follows

$$f_{i,PAR}(z) = \frac{1}{PAR\ (z)^{*}\sigma_i\sqrt{2\pi}}\ exp(-\frac{(\ ln(PAR\ (z))-\mu_i)^2}{2\sigma_i^2})$$

where $\sigma_i^2 = ln(1 + \frac{S_i^2}{m_i^2})$ and $\mu_i = ln\left(m_i\right) - \frac{\sigma_i^2}{2}$ . $m_i$ and $S_i$ being the desired mean and variance values of the distribution, which are both assumed to be community-dependent. *PAR(z)* is the intensity of the incident light at depth level z. The *mode* value representing the PAR intensity corresponding to the global maximum of the distribution is written as

$$mode_i = exp(\mu_i - \sigma_i^2)$$

This means that when $PAR(z) = mode_i$ the functional response $f_{i,PAR}(z) = f_{i,mode_I}(z)$ will reach its maximum value. Therefore, to ensure that the score values range between 0 and 1, the functional response $f_{i,PAR}(z)$ was normalized with respect to $f_{i,mode_I}(z)$ leading to the following light preference function

$$pref_{i,PAR}(z) = \frac{mode_i}{PAR(z)}\ exp(-\frac{(ln(mode_i)-\mu_i)^2-(ln(PAR(z))-\mu_i)^2}{2\sigma_i^2}) \qquad (S1)$$

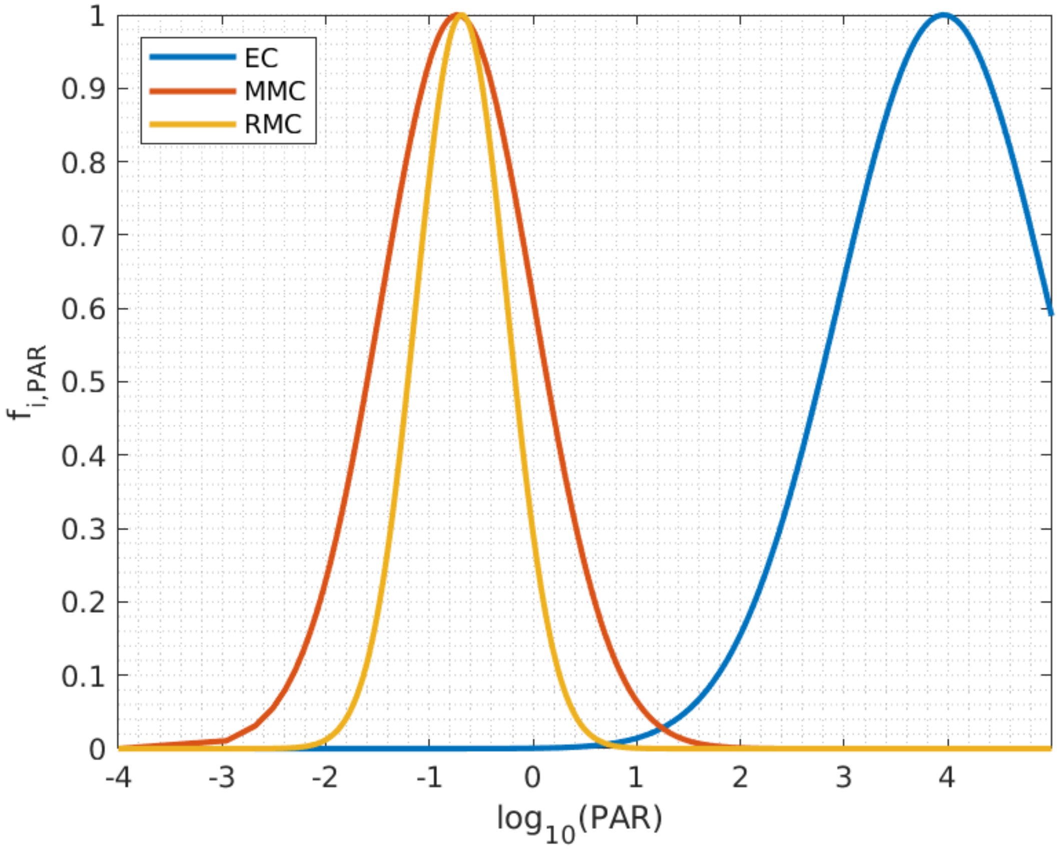


**Fig.S1** : Functional response to underwater light intensity estimated from equation S1 and shown for the three pelagic communities, Epipelagic community (EC), Migratory mesopelagic community (MMC) and Resident mesopelagic community (RMC). The parameters of the equation S1 are estimated by calibration.

##### Functional Response to Dissolved Oxygen

Oxygen is very important for the metabolism and survival of marine organisms. Therefore, oxygen concentrations in seawater have been found to correlate with the daytime vertical positions of the mesopelagic communities especially in the oxygen-poor waters (7–10) . The authors hypothesized that hypoxic boundaries in the ocean could form a sharp barrier preventing most migratory organisms from descending deeper into the water column and could, therefore, limit the maximum depth available as suitable habitat for the mesopelagic organisms. To account for this effect, the response of each generic community *com* to the dissolved oxygen concentration was expressed as a logistic function

$$Pref_{i,oxy}(z) = \frac{1}{1 + exp(a_{i,oxy}(b_{i,oxy} - [O_2(z)]))} \qquad (S2)$$

where $[O_2(z)]$ is the concentration of the dissolved oxygen at a depth level z of the water column, $a_{i,\,oxy}$ is the steepness of the curve and $b_{i,\,oxy}$ is the half-saturation coefficient for oxygen limitation for the generic community $i$.

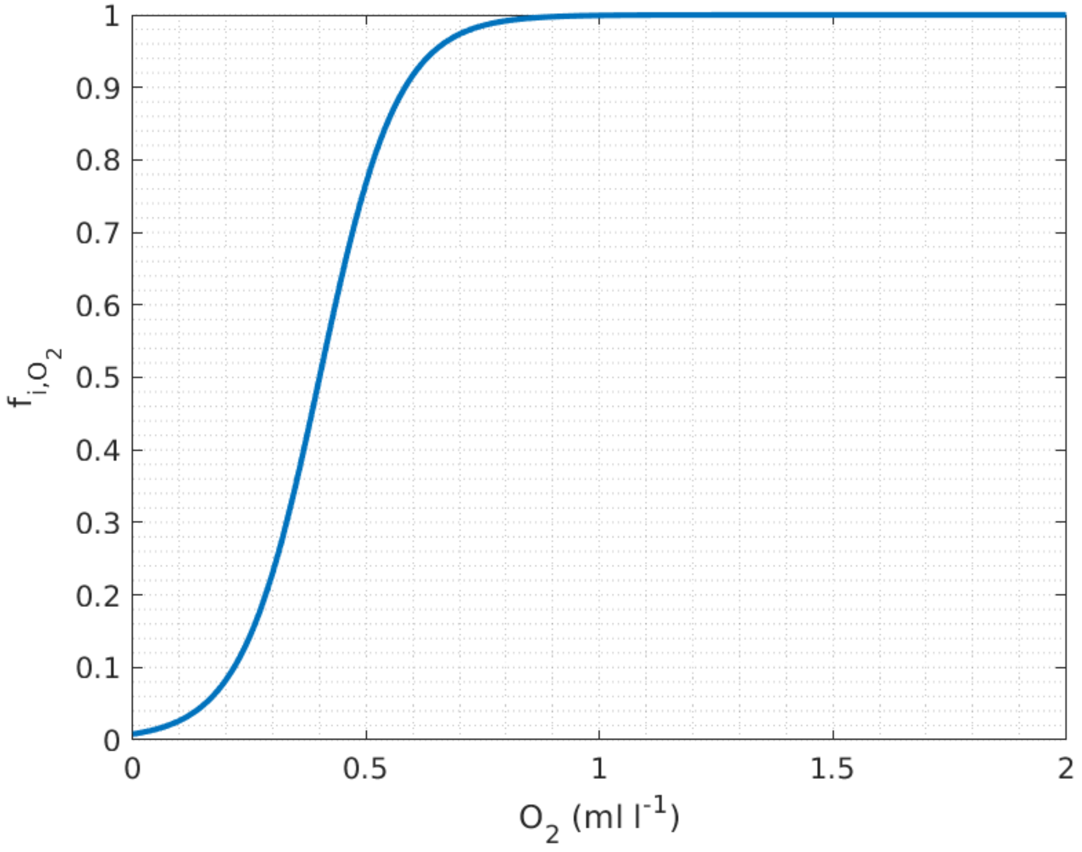


**Fig.S2** : Functional response to dissolved oxygen concentrations in the water estimated from equation S2 shown for the Migratory mesopelagic community. The parameters of the equation S2 are estimated by calibration.

**Functional response to water Temperature**

The physiological and metabolic processes of marine organisms depend on temperature. It is therefore an important factor structuring the vertical distribution of marine communities, in particular the epipelagic community.

The functional response to the water temperature is represented by a normal distribution

$$Pref_{i,T}(z) = exp\left( -\frac{1}{2}\left(\frac{(T_{cor}(z) /- T_{cor}(0))-}{\delta_{Tcor}}\right)^2 \right) \qquad (S3)$$

where, $T_{cor}(z) = exp(\frac{T_a}{T_{ref}} - \frac{T_a}{T(z)})$ describes the changes of any physiological rate with temperature, $T_a$ being the Arrhenius temperature, $T_{ref}$ is a reference temperature and $T(z)$ the water temperature at a vertical level $z$.

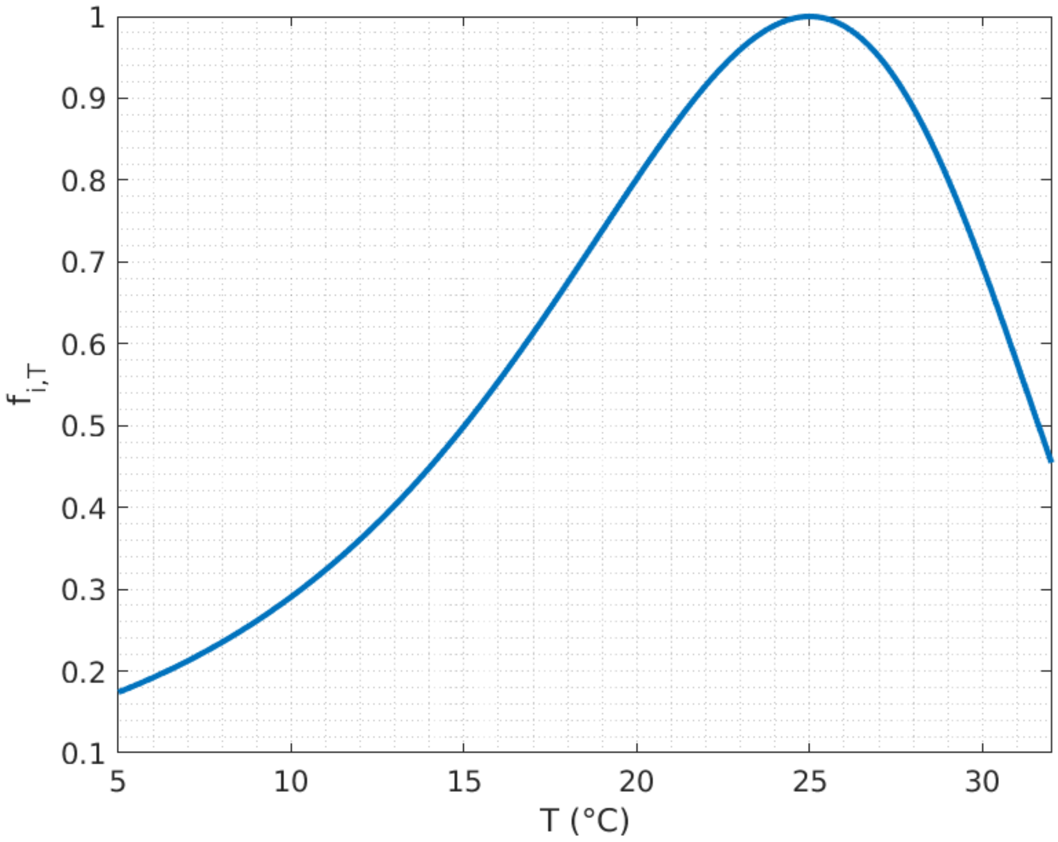


**Fig.S3** : Functional response to water temperature estimated from equation S3. The parameters of the equation S3 are estimated by calibration.

#### 2) Relative biomasses

To calculate the relative biomasses for the generic communities represented in the model, we used the echosounder observations collected during the Malaspina cruise (see main text). The relative proportions of the three generic communities (EC, MMC, RMC) are calculated from the acoustic profiles as follow:

- For the epipelagic community, it is assumed to be the relative proportion of the daytime acoustic backscatter within the upper 200m of the ocean.

  $\alpha_{EC}(s) = \frac{\sum_{z_1}^{z_{200}} A_d(z,s)}{\sum_{z_{200}}^{z_{max}} A_d(z,s)}$ , where *s* refers to the station and z to the depth level. $A_d$ is the daytime acoustic profile. $z_1$, $z_{200}$, and $z_{max}$ refer to the depth levels corresponding to 0, 200 and 1000 m depth, respectively.

- For the mesopelagic communities (MMC and RMC), both daytime and night-time acoustic vertical profiles are used:

  $\alpha_{RMC}(s) = (1 - \alpha_{EC}(s)) * \frac{\sum_{z_{200}}^{z_{max}} A_n(z,s)}{\sum_{z_{200}}^{z_{max}} A_d(z,s)}$ , where $A_n$ is the night-time acoustic profile

  $\alpha_{MMC}(s) = 1 - \alpha_{EC}(s) - \alpha_{RMC}(s)$

In the case of the model configuration with shallower and deeper mesopelagic subcommunities, the relative proportion of each subcommunity is calculated as follows

$$\alpha_{RMC_S}(s) = \beta_{RMC}(s) * \alpha_{RMC}(s)$$

$$\alpha_{RMC_D}(s) = (1 - \beta_{RMC}(s)) * \alpha_{RMC}(s)$$

$$\alpha_{MMC_S}(s) = \beta_{MMC}(s) * \alpha_{MMC}(s)$$

$$\alpha_{MMC_D}(s) = (1 - \beta_{MMC}(s)) * \alpha_{MMC}(s)$$

where $\beta_{RMC}$ and $\beta_{MMC}$ are spatially dependent parameters representing the proportions of shallower resident and migratory subcommunities, respectively.

#### 3) Model Calibration

We calibrated the model using the AD Model Builder(11) . ADMB calculates the exact derivatives of the objective function with the automatic differentiation technique, which makes it possible to fit models with a large number of parameters in an efficient and reliable manner.

In this study, we have two kinds of parameters: (i) Spatially independent parameters that are constant everywhere; and (ii) Spatially dependent parameters which are different from one station to another.

For a better performance of the calibration, we used a multi-phase approach:

- In the initial phase, the minimization is carried out over a subset of the community-dependent parameters while keeping constant the spatially dependent parameters (relative community biomasses).
- The relative community biomasses are included into the minimization in a number of phases equivalent to the number of stations (one station at each phase).

For simplicity, we assume that the size distribution of marine organisms follows a Sheldon spectrum(12). Therefore, the abundance ratio $\kappa(L)$ between a given size-classe $L$ with respect to the smaller size-class $L_1$ can be expressed as follows(13)

$$\kappa(L) = \frac{N(L)}{N(L_1)} = \left(\frac{L}{L_1}\right)^{-4}$$

The correction factor implemented in the model calibration to account for the nonlinear distortion of the acoustic signal with depth and organisms size is calculated as follows:

$$\phi(L, z) = \kappa(L) * \sigma_{bs}(L, z)$$

where, $\sigma_{bs}(L, z)$ is the expected backscattering cross-section of one insonified fish belonging to the size-class $L$ at the depth level $z$. It was estimated from Monte-carlo simulations using an acoustic model (see below).

#### 3.1 Acoustic Model

In this study, we assume that acoustic backscattering intensity produced at 38 kHtz frequency by gas-bladdered fish and siphonophores is widely dominant (~100% of contribution) compared to that produced by other marine organisms such as gelatinous species, copepods, squid, other fish and crustacean species, as highlighted in several previous studies (14–17) . For this purpose, we attribute the observed backscattering intensity only to fish. This is reasonable given that, at a global scale, siphonophores are much less abundant than fish organisms especially in the mesopelagic zone (16) . Moreover, the distribution of the gas-bladerred fish individuals in each pelagic community is assumed homogeneous throughout the water column.

The acoustic model presented here and its parameterization are mainly based on the studies by *Love (1978)*(18) , *Scoulding et al. (2015)*(19) and *Proud et al.(2019)*(16) .

Given that swimbladders contribute much more than fish flesh to backscattered acoustic intensity, only swimbladder is used in calculating the backscattering cross-section. The shape of the swimbladder is assumed to be prolate spheroid, and its equivalent spherical radius $a_{esr}$ is calculated as follows

$$a_{esr}(L) \;=\; \left(\frac{3\,V_{swb}(L)}{4\pi}\right)^{1/3}$$

where the volume of the fish swimbladder $V_{swb}$ is assumed to be proportional to the volume of the fish body $V_f$ such as : $V_{swb}(L) \;=\; P_{swb}\, V_f(L)$ , with $P_{swb}$ being the proportion of the swimbladder with respect to the total volume of the fish individual. To simplify, the shape of the fish body is assumed to be a prolate spheroid, its volume is calculated as a function of the fish length (L) as follows

$$V_f(L) \;= \frac{\pi}{6\,\alpha^2}\, L^3$$

The resonance frequency of the fish swimbladder $f_{res}$ is related to $a_{esr}$ by the following equation

$$f_{res}(L, z) \;=\; \frac{1}{2\pi\, a_{esr}(L)}\sqrt{\frac{3\,\gamma_a P(z) + 4\,\mu_r}{\rho_w}}$$

where $\gamma_a$ is the ratio of the specific heat for air, *P(z)* is the ambient pressure at depth z, $\mu_r$ is the real part of the rigidity of fish flesh, and $\rho_w$ is the density of water.

Acoustic backscattering cross-section at 38kHz incident frequency can be estimated using the following equation

$$\sigma_{bs}(L, z) \;= \frac{a_{esr}(L,z)^2 \left(\frac{\rho_w}{\rho_f}\right)^2}{\left(\left(\frac{f_{res}(L,z)}{f_{38}}\right)^2 - 1\right)^2 \;+\; d_f^2}$$

with $d_f$ being the damping factor including radiation, viscous and thermal processes, calculated as in Scoulding et al. (2015)(19) .

#### 3.2 Monte Carlo simulations

To take into account the uncertainty of some model parameters (), we carried out Monte Carlo simulations using the bootstrap sampling method. We run 1000 simulations, each time with a different set of parameters selected randomly from a range of variability of each parameter (Table S1). Therefore, we estimated 1000 vertical profile (0-1000m depth) of $\sigma_{bs}$ for each of the size classes considered in this study. For simplicity, we used for each size-class the mean vertical profile of $\sigma_{bs}$ (averaged over the 1000 obtained profiles) (Fig.S4).

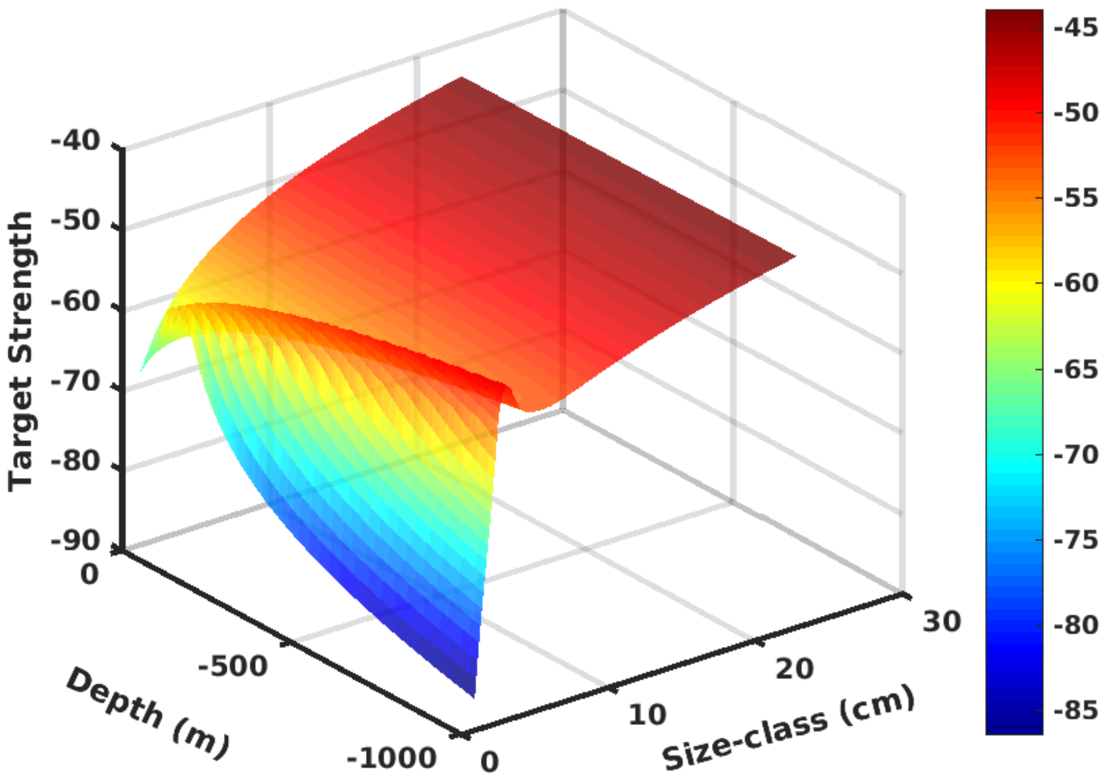


**Fig.S4** : Mean values of Target strength ($TS = 10\ log_{10}(\sigma_{bs})$ ) displayed as a function of water depth and fish length.

**Table S1**: Parameters of the acoustic model

| Parameter | Description | value (std) | remark |
|---|---|---|---|
| $f_{38}$ | incident acoustic frequency | 38000 | constant |
| $\gamma_a$ | Ratio of the specific heat for air | 1.4 | constant |

| | | | |
|---|---|---|---|
| $\rho_w$ | Density of seawater | 1028 | constant |
| $\rho_f$ | Density of fish flesh | 1050 | constant |
| $\mu_r$ | Real part of the rigidity of fish flesh | $10^5(10^4)$ | normal distribution |
| $\alpha$ | Aspect ratio of fish body | 8(2) | normal distribution |
| $P_{swb}$ | Swimbladder volume as a proportion of fish volume | 0.01(0.002) | normal distribution |

#### 4) Global application of the model

The model was applied globally to estimate the vertical distributions of the 5 pelagic communities. The spatial distributions of the predicted weighted mean depths (WMDs) are shown in Fig.S4 for the five generic communities considered in this study (EC, $MMC_S$ , $MMC_D$ , $RMC_S$ , $RMC_D$ ). The results show deeper distributions (higher WMD values) in the subtropical and subpolar gyres, and shallower distributions (lower WMD values) in the tropical and polar regions especially in the poor-oxygen waters such as the oxygen minimum zones of the Eastern Pacific, Arabian sea and Northern Indian ocean. Overall, these predicted distribution patterns are very consistent with the spatial distributions of the mean vertical positions of pelagic organisms estimated from acoustic observations at a global scale (7)

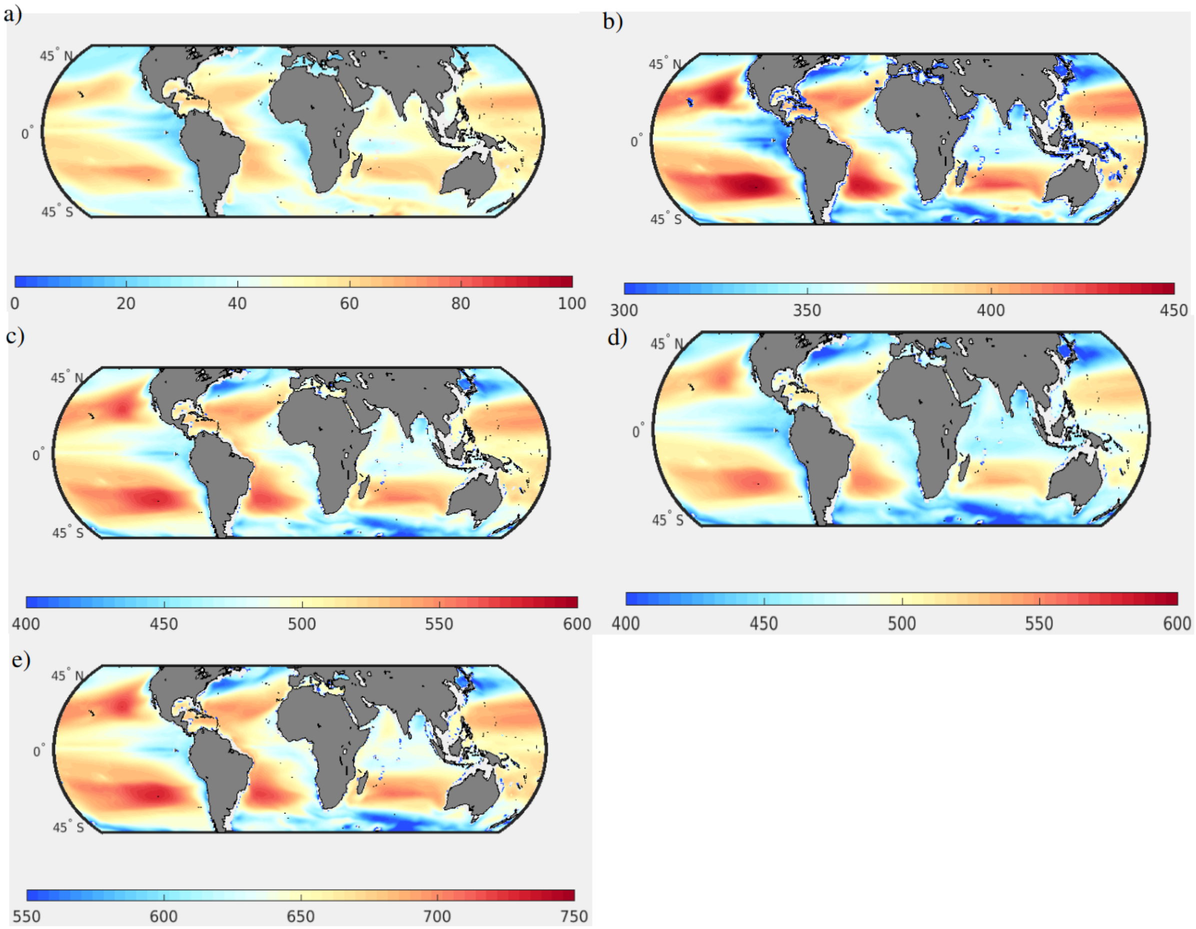


**Fig.S4** Spatial distribution of the annual mean of WMD predicted for the five generic communities considered in the model: a) EC , b) $MMC_S$ , c) $MMC_D$, d) $RMC_S$ (e) $RMC_D$

#### 5) Multi-layer DSLs in specific regions of the global ocean

A detailed inspection of the Malaspina echograms reveals that DSLs are not always homogenous and sometimes characterized by two (or more) distinct layers often composed of migrating and resident organisms as shown in Fig. S5 for two different geographic regions.

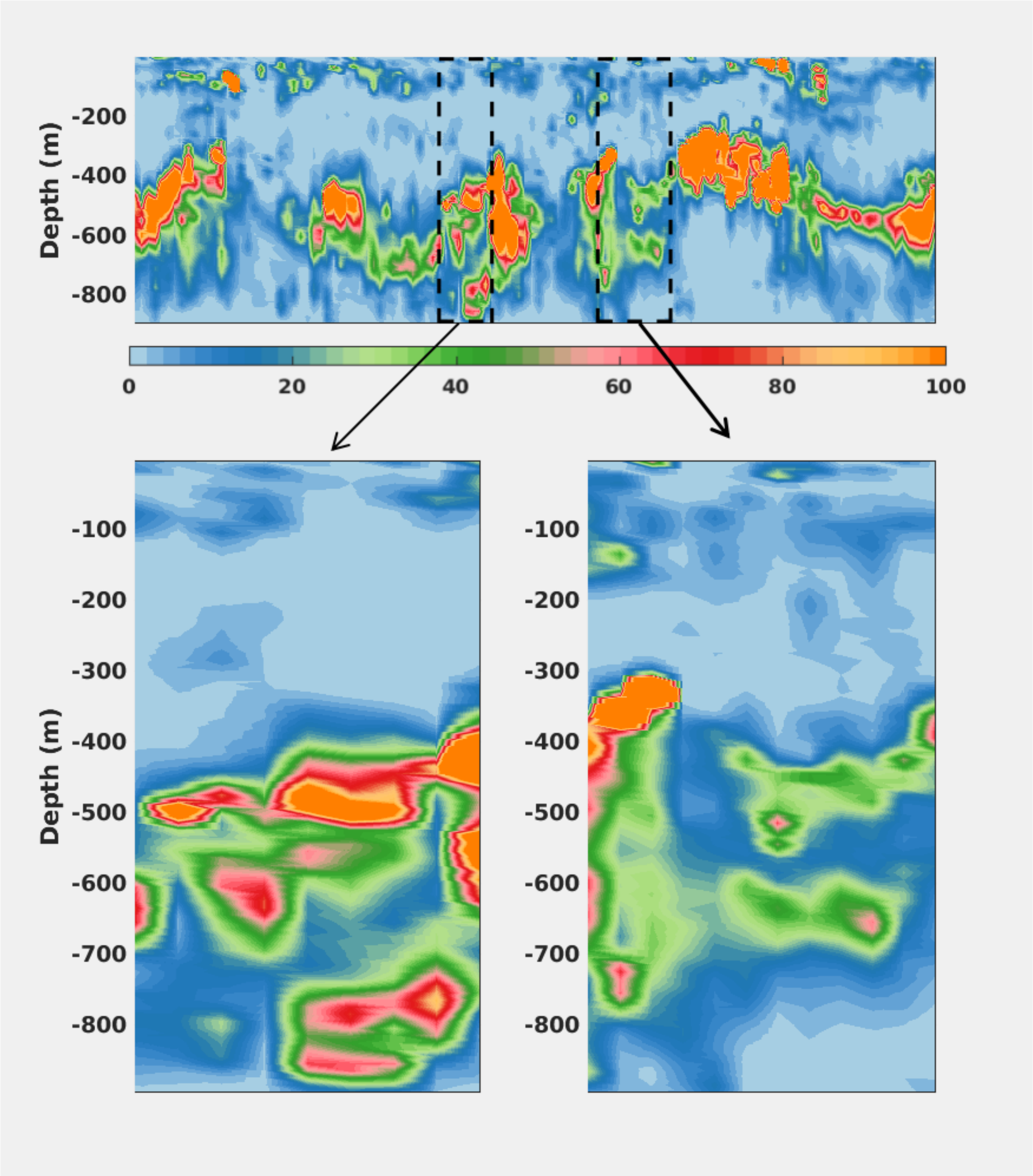


**Fig. S5** : Daytime echogram at 38 kHz along the Malaspina cruise track, showing two geographic regions with multiple DSLs.

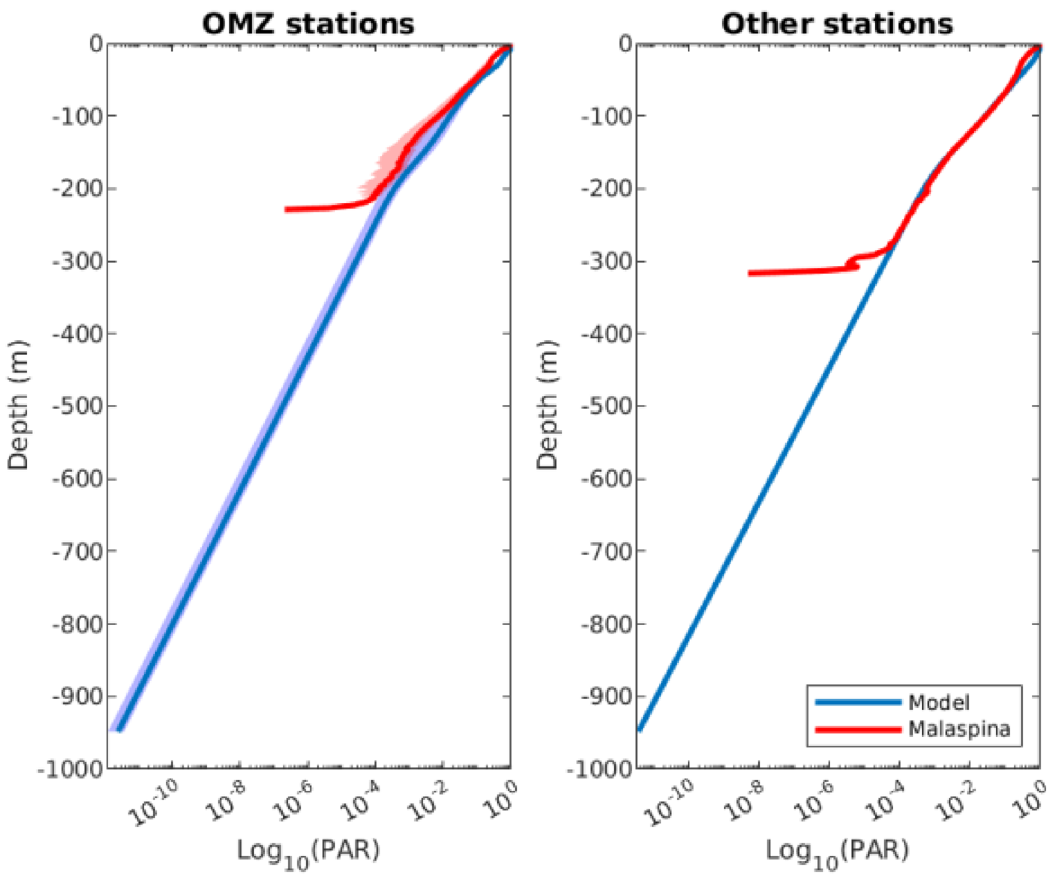


**Fig. S6 :** Mean vertical distribution of PAR normalized with respect to the maximum value (at the surface), as provided by Malaspina observations (red) and the PISCES model (blue) averaged at OMZ stations (left) and non-OMZ stations (right). The shaded areas correspond to the confidence interval.

### References


1. E. Norheim, T. Klevjer, D. Aksnes, Evidence for light-controlled migration amplitude of a sound scattering layer in the Norwegian Sea. *Mar. Ecol. Prog. Ser.* **551**, 45–52 (2016).

2. A. Røstad, S. Kaartvedt, D. L. Aksnes, Light comfort zones of mesopelagic acoustic scattering layers in two contrasting optical environments. *Deep Sea Res. Part Oceanogr. Res. Pap.* **113**, 1–6 (2016).

3. D. L. Aksnes, *et al.*, Light penetration structures the deep acoustic scattering layers in the global ocean. *Sci. Adv.* **3**, e1602468 (2017).

4. S. Kaartvedt, T. J. Langbehn, D. L. Aksnes, Enlightening the ocean's twilight zone. *Ices J. Mar. Sci.* **76**, 803–812 (2019).

5. W. J. O'Brien, The Predator-Prey Interaction of Planktivorous Fish and Zooplankton: Recent research with planktivorous fish and their zooplankton prey shows the evolutionary thrust and parry of the predator-prey relationship. *Am. Sci.* **67**, 572–581 (1979).

6. D. L. Aksnes, J. Nejstgaard, E. Sædberg, T. Sørnes, Optical control of fish and zooplankton populations. *Limnol. Oceanogr.* **49**, 233–238 (2004).

7. D. Bianchi, E. D. Galbraith, D. A. Carozza, K. A. S. Mislan, C. A. Stock, Intensification of open-ocean oxygen depletion by vertically migrating animals. *Nat. Geosci.* **6**, 545–548 (2013).

8. A. Bertrand, M. Ballon, A. Chaigneau, Acoustic Observation of Living Organisms Reveals the Upper Limit of the Oxygen Minimum Zone. *Plos One* **5**, e10330 (2010).

9. A. N. Netburn, J. Anthony Koslow, Dissolved oxygen as a constraint on daytime deep scattering layer depth in the southern California current ecosystem. *Deep Sea Res. Part Oceanogr. Res. Pap.* **104**, 149–158 (2015).

10. T. A. Klevjer, *et al.*, Large scale patterns in vertical distribution and behaviour of mesopelagic scattering layers. *Sci. Rep.* **6**, 19873 (2016).

11. D. A. Fournier, *et al.*, AD Model Builder: using automatic differentiation for statistical inference of highly parameterized complex nonlinear models. *Optim. Methods Softw.* **27**, 233–249 (2012).

12. R. W. Sheldon, A. Prakash, W. H. Sutcliffe, The Size Distribution of Particles in the Ocean1. *Limnol. Oceanogr.* **17**, 327–340 (1972).

13. J. Guiet, J.-C. Poggiale, O. Maury, Modelling the community size-spectrum: recent developments and new directions. *Ecol. Model.* **337**, 4–14 (2016).

14. A. C. Lavery, D. Chu, J. N. Moum, Measurements of acoustic scattering from zooplankton and oceanic microstructure using a broadband echosounder. *ICES J. Mar. Sci.* **67**, 379–394 (2010).

15. P. C. Davison, J. A. Koslow, R. J. Kloser, Acoustic biomass estimation of mesopelagic fish: backscattering from individuals, populations, and communities. *ICES J. Mar. Sci.*

**72**, 1413–1424 (2015).

16. R. Proud, N. O. Handegard, R. J. Kloser, M. J. Cox, A. S. Brierley, From siphonophores to deep scattering layers: uncertainty ranges for the estimation of global mesopelagic fish biomass. *ICES J. Mar. Sci.* **76**, 718–733 (2019).

17. R. J. Kloser, T. E. Ryan, G. Keith, L. Gershwin, Deep-scattering layer, gas-bladder density, and size estimates using a two-frequency acoustic and optical probe. *ICES J. Mar. Sci.* **73**, 2037–2048 (2016).

18. R. H. Love, Resonant acoustic scattering by swimbladder-bearing fish [a)]. *J. Acoust. Soc. Am.* **64**, 571–580 (1978).

19. B. Scoulding, D. Chu, E. Ona, Paul. G. Fernandes, Target strengths of two abundant mesopelagic fish species. *J. Acoust. Soc. Am.* **137**, 989–1000 (2015).
